# CORK1, a LRR-Malectin Receptor Kinase for Cellooligomer Perception in *Arabidopsis thaliana*

**DOI:** 10.1101/2022.04.29.490029

**Authors:** Yu-Heng Tseng, Sandra S. Scholz, Judith Fliegmann, Thomas Krüger, Akanksha Gandhi, Olaf Kniemeyer, Axel A. Brakhage, Ralf Oelmüller

## Abstract

Cell wall integrity (CWI) maintenance is central for plant cells. Mechanical or chemical distortions, pH changes, or breakdown products of cell wall polysaccharides activate plasma membrane-localized receptors and induce appropriate downstream responses. Microbial interactions alter or destroy the structure of the plant cell wall, connecting CWI maintenance to immune responses. Cellulose is the major polysaccharide in the primary and secondary cell wall. Its breakdown generates short-chain cellooligomers which induce Ca^2+^-dependent CWI responses. We have shown here that these responses require the malectin domain-containing CELLOOLIGOMER-RECEPTOR KINASE 1 (CORK1) in *Arabidopsis*. CORK1 is required for cellooligomer-induced cytoplasmic Ca^2+^ elevation, reactive oxygen species (ROS) production, mitogen associated protein kinase (MAPK) activation, cellulose synthase phosphorylation, and the regulation of CWI-related genes including those involved in biosynthesis of cell wall material, secondary metabolites and tryptophan. Phosphoproteome analyses identified early targets involved in signaling, cellulose synthesis, the endoplasmatic reticulum/Golgi secretory pathway, cell wall repair and immune responses. Two conserved phenylalanine residues in the malectin domain are crucial for CORK1 function. We propose that cellulose breakdown products bind to the malectin domain in CORK1, indicating its role as a novel receptor kinase for CWI maintenance.

## Introduction

The plant cell wall is mainly composed of the polysaccharide polymers cellulose, hemicellulose and pectin. Cellulose accounts for more than 30% of the total plant cell wall material (Lodish et al., 2000), and consists of β-(1,4)-bound D-glucose moieties, which form unbranched fibers with a paracrystaline structure. Hemicellulose is made of xylans, xyloglucans, mannans, glucomannans, and β-(1,3;1-4)-glucans. The backbone for xylans, xyloglucans and mannans is made of β-(1,4) linked monomer residues, while β-(1,3;1-4)-glucans contain β-(1-4)-linked glucose interleaved with β-(1,3) linkages. Unlike cellulose, hemicellulose has short branches and its amorphous structure is easily accessible to hydrolases (Scheller and Ulvskov, 2010; Gibson, 2012). Pectins are categorized into unbranched homogalacturonan (HG), branching rhamnose (Rha)- containing rhamnogalacturonan I (RG-I), and rhamnogalacturonan II (RG-II) with complex composition (Mohnen, 2008; Atmodjo et al., 2013). While HG is a monopolymer of α-(1,4) galacturonic acid (GalA), RG-I has a disaccharide unit backbone of α-D-GalA-(1,2)- α-L-Rha, and RG-II possesses GalA linked with various sugars (Ridley et al., 2001; Harholt et al., 2010).

Fragments of these cell wall polysaccharides have been shown to act as damage-associated molecular patterns (DAMPs; Gust et al., 2017). Cellulose breakdown products, cellooligomers (COMs), trigger calcium influx, ROS production, MAPK phosphorylation, and defense-related gene expression, which eventually lead to higher pathogen resistance (Hu et al., 2004; Galletti et al., 2011; Souza et al., 2017; Claverie et al., 2018; Johnson et al., 2018; Mélida et al., 2020; Yang et al., 2021a). COMs with 2-7 glucose moieties induce cytoplasmic calcium ([Ca^2+^]_cyt_) elevation. The amplitude of the response depends on the length of the oligomer, and cellotriose (CT) was the most active COM in *Arabidopsis thaliana* (Johnson et al., 2018). The defense responses induced by COMs are relatively mild when compared to those induced by the pathogen-associated molecular patterns (PAMPs) chitin or flg22 (Souza et al., 2017; Claverie et al., 2018; Johnson et al., 2018; Rebaque et al., 2021). However, in combination with chitin, flg22 or oligogalacturonic acid (OG), synergistic effects on calcium influx, ROS production, and MAPK phosphorylation indicate crosstalk between COM and PAMP responses (Souza et al., 2017; Johnson et al., 2018).

Plants rely on an array of membrane-associated pattern recognition receptors (PRRs) to recognize breakdown products of its cell wall. The wall-associated kinase 1 (WAK1) is activated by the pectin fragments OGs (He et al., 1996; Brutus et al., 2010). FERONIA, a membrane-localized receptor-like kinase with a malectin-like domain, is involved in monitoring cell wall integrity (CWI), pollen tube development, plant growth and perception of RALF peptides (Escobar-Restrepo et al., 2007; Haruta et al., 2014). Its extracellular region has been shown to interact with pectin (Feng et al., 2018; Tang et al., 2021). In rice, two species of mixed-linked β-1,3/1,4-glucans (MLGs) from hemicellulose, namely 3^1^-β-D-cellobiosyl-glucose and 3^1^-β-D-cellotriosyl-glucose, bind to OsCERK1, and induce the dimerization of OsCERK1 and the chitin receptor OsCEBiP (Yang et al., 2021a). However, up to date, no receptor has been reported to perceive β-1,4 glucans. Here, we show that CORK1 (<u>C</u>ello<u>O</u>ligomer <u>R</u>eceptor <u>K</u>inase 1) specifically recognizes

COMs in *A. thaliana*. CORK1 is a functional LRR-malectin receptor kinase. Upon COM treatment, *CORK1* mutants are impaired in [Ca^2+^]_cyt_ elevation, ROS production, regulation of genes involved in CWI maintenance and immune responses, including *WRKY30*/*WRKY40*. Two conserved phenylalanine residues in the malectin domain are crucial for COM-induced responses in *A. thaliana*. Phosphoproteome and transcriptome data identified protein and gene targets of the novel COM/CORK1 pathway and have shed light on the role of COMs in CWI maintenance.

## Materials and Methods

### Growth medium and conditions for seedlings

*A. thaliana* seeds were surface-sterilized for 8 minutes in sterilization solution containing lauryl sarcosine (1%) and Clorix cleaner (23%). Surface-sterilized seeds were washed with sterilized water 8 times and placed on Petri dishes with MS medium supplemented with 0.3% gelrite (Murashige and Skoog, 1962). After cold treatment at 4°C for 48 hours, plates were incubated at 22°C under long day conditions (16 hours light/ 8 hours dark; 80 μmol m^−2^ s^−1^).

Wild-type (ecotype Columbia-0), the aequorin-containing wild-type [pMAQ2] line (AeqWT; Knight et al., 1991), and EMS (ethyl methanesulfonate)-induced mutant lines in AeqWT background (Johnson et al., 2018) were used in this study. In addition, two T-DNA insertion lines, *cork1-1* (N671776; SALK_099436C) and *cork1-2* (N674063; SALK_021490C), were obtained from Nottingham *Arabidopsis* Stock Centre (NASC). Homozygous seedlings of these insertion lines were crossed to the AeqWT. The corresponding segregated wild-type (SWT) and homozygous (HO) seedlings from the F3 generation were used for experiments.

### EMS mutagenesis of *A. thaliana* seeds

2.5 g of AeqWT seeds were used for mutagenesis. According to Kim *et al*. (2006), seeds were soaked in 40 ml of 100 mM phosphate buffer (pH = 7.5) for 10 hours at 4°C. The next day, the buffer was replaced and EMS (Sigma-Aldrich, Germany) was added to a final concentration of 0.2 %. The mixture was incubated at room temperature in a hood over night with gentle stirring. The seeds were washed twice in 40 ml of 100 mM sodium thiosulphate for 15 min to destroy the remaining EMS, followed by 18 wash steps with water (Leyser and Furner, 1993). Freshly mutagenized seeds were directly separated in different Eppendorf tubes, surface-sterilized and germinated as described above. Three-week old plants were transferred to soil to obtain the seeds of the individual mother plants.

### Whole Genome Sequencing and SNP analysis

After screening ∼100 independent EMS lines, we found a COM non-responsive mutant, named here as EMS71. From the F2 generation of the back-cross between EMS71 and AeqWT, two pools of seedlings were sorted out. One pool consisted of CT responsive individuals, while the other contained non-responsive individuals (50 seedlings in each pool). Whole genome sequencing of the two pools was performed on Illumina sequencing platform (PE150; Novogene Co., UK). The reads from both pools were mapped separately against the TAIR10 reference genome using Samtools v. 1.8 (Li et al., 2009). SIMPLE v. 1.8.1 (Wachsman et al., 2017) analysis was implemented to filter out single-nucleotide polymorphisms (SNPs) which appeared 90-100 % in the non-responsive population, while less than 70% in the responsive population. Putative candidate genes were selected based on whether the SNP causes non-synonymous mutation, or affects mRNA processing (e.g. mRNA splicing). SNP sites of candidate genes were confirmed in 3 different individuals from the EMS71 line. As a result, two candidate genes, *ARF1* (At1g59750) and *CORK1* (AT1G56145), were selected for further confirmation as described in the Result section.

### Transcriptome analysis

16 roots of SWT or HO from *cork1-2* mutant line crossed to AeqWT were treated with 1 mL of either water or 10 µM CT for 1 hour. Total RNA was extracted and purified as described, and sent to Novogene Co. (UK) for sequencing with Illumina NovaSeq instrument (poly-A enrichment; PE150). The raw reads were aligned to *Arabidopsis* TAIR10 reference genome using STAR v. 2.7.10a (Dobin et al., 2013). The aligned bam files were analyzed with featureCounts v. 2.0.1 (Liao et al., 2014), and the count table for all samples were analyzed with DESeq2 v. 1.34.0 (Love et al., 2014). GO enrichment and KEGG pathway analysis was performed on PANTHER (Mi et al., 2021) and KEGG PATHWAY (Kanehisa and Goto, 2000) database, respectively. Significantly regulated genes were defined with the criteria: |log2 fold change| ≥ 1.33 and adjusted p-value < 0.05. The adjusted p-value was calculated by DESeq2 using the built-in Benjamini and Hochberg method. The default FDR cutoff value was set as 0.1.

### Phosphoproteomic analysis

#### Sample collection

300 roots of SWT or HO from the *cork1-2* mutant line were collected at 0 minute, or after treatment with either water or 10 µM CT for 5 or 15 minutes. Samples were immediately frozen in liquid nitrogen until further analysis.

#### In-solution digest

Tissues were disrupted by using mortar and pestle with liquid nitrogen. Debris were homogenized in lysis buffer (1% (w/v) SDS, 150 mM NaCl, 100 mM TEAB (triethyl ammonium bicarbonate)), one tablet each of cOmplete Ultra Protease Inhibitor Cocktail and PhosSTOP). After addition of 0.5 µl Benzonase nuclease (250 U/μl) the samples were incubated at 37°C in a water bath sonicator for 30 min. Proteins were separated from unsolubilized debris by centrifugation (15 min, 18000 × *g*). Each 1.5 mg of total protein per sample was diluted with 100 mM TEAB to gain a final volume of 1.5 ml. Subsequently, cysteine thiols were reduced and carbamidomethylated in one step for 30 min at 70°C by addition of 30 µL of 500 mM TCEP (tris(2-carboxyethyl)phosphine) and 30 µl of 625 mM 2-chloroacetamide (CAA). The samples were further cleaned up by methanol-chloroform-water precipitation using the protocol of Wessel and Flügge (1984). Protein precipitates were resolubilized in 5% trifluoroethanol of aqueous 100 mM TEAB and digested overnight (18 hours) with a Trypsin+LysC mixture (Promega) at a protein to protease ratio of 25:1. Each sample was divided in 3 × 0.5 mg used for the phosphopeptide enrichment and 150 µg initial protein used for the reference proteome analysis. Samples were evaporated in a SpeedVac. The reference proteome sample was resolubilized in 30 µL of 0.05% TFA in H_2_O/ACN 98/2 (v/v) filtered through 10 kDa MWCO PES membrane spin filters (VWR). The filtrate was transferred to HPLC vials and injected into the LC-MS/MS instrument.

#### Phosphopeptide enrichment

Phosphopeptides were enriched by using TiO_2_+ZrO_2_ TopTips (Glygen Corp., Columbia, MD, USA). TopTips were loaded with 0.5 mg protein isolate using 3 TopTips per biological replicate after equilibration with 200 µl Load and Wash Solution 1, LWS1 (1% trifluoroacetic acid (TFA), 20% lactic acid, 25% acetonitrile (ACN), 54% H_2_O). TopTips were centrifuged at 1500 rpm (∼200 × *g*) for 5 min at room temperature. After washing with 200 µl LWS1, the TiO_2_/ZrO_2_ resin was washed with 25% ACN and subsequently the phosphopeptides were eluted with 200 µl NH_3_·H_2_O (NH_4_OH), pH 12. The alkaline solution was immediately evaporated using a SpeedVac. The phosphoproteome samples were resolubilized in 50 µL of 0.05% TFA in H_2_O/ACN 98/2 (v/v) filtered through 10 kDa MWCO PES membrane spin filters (VWR). The filtrate was also transferred to HPLC vials and injected into the LC-MS/MS instrument.

#### LC-MS/MS analysis

Each sample was measured in duplicate (2 analytical replicates of 3 biological replicates of a reference proteome fraction and a phosphoproteome fraction). LC-MS/MS analysis was performed on an Ultimate 3000 nano RSLC system connected to a QExactive HF mass spectrometer (both Thermo Fisher Scientific, Waltham, MA, USA). Peptide trapping for 5 min on an Acclaim Pep Map 100 column (2 cm × 75 µm, 3 µm) at 5 µL/min was followed by separation on an analytical Acclaim Pep Map RSLC nano column (50 cm × 75 µm, 2µm). Mobile phase gradient elution of eluent A (0.1% (v/v) formic acid in water) mixed with eluent B (0.1% (v/v) formic acid in 90/10 acetonitrile/water) was performed using the following gradient: 0-5 min at 4% B, 30 min at 7% B, 60 min at 10% B, 100 min at 15% B, 140 min at 25% B, 180 min at 45% B, 200 min at 65% B, 210-215 min at 96% B, 215.1-240 min at 4% B. Positively charged ions were generated at spray voltage of 2.2 kV using a stainless steel emitter attached to the Nanospray Flex Ion Source (Thermo Fisher Scientific). The quadrupole/orbitrap instrument was operated in Full MS / data-dependent MS2 Top15 mode. Precursor ions were monitored at m/z 300-1500 at a resolution of 120,000 FWHM (full width at half maximum) using a maximum injection time (ITmax) of 120 ms and an AGC (automatic gain control) target of 3 × 10^6^. Precursor ions with a charge state of z=2-5 were filtered at an isolation width of *m/z* 1.6 amu for further HCD fragmentation at 27% normalized collision energy (NCE). MS2 ions were scanned at 15,000 FWHM (ITmax=100 ms, AGC= 2 × 10^5^) using a fixed first mass of *m/z* 120 amu. Dynamic exclusion of precursor ions was set to 30 s. The LC-MS/MS instrument was controlled by Chromeleon 7.2, QExactive HF Tune 2.8 and Xcalibur 4.0 software.

#### Protein database search

Tandem mass spectra were searched against the UniProt database (2022/01/06; https://www.uniprot.org/proteomes/UP000006548) of *Arabidopsis thaliana* using Proteome Discoverer (PD) 2.4 (Thermo) and the Sequest HT algorithm. Two missed cleavages were allowed for the tryptic digestion. The precursor mass tolerance was set to 10 ppm and the fragment mass tolerance was set to 0.02 Da. Modifications were defined as dynamic Met oxidation, phosphorylation of Ser, Thr, and Tyr, protein N-term acetylation with and without Met-loss as well as static Cys carbamidomethylation. A strict false discovery rate (FDR) < 1% (peptide and protein level) and an X_corr_ score > 4 was required for positive protein hits. The Percolator node of PD2.4 and a reverse decoy database was used for qvalue validation of spectral matches. Only rank 1 proteins and peptides of the top scored proteins were counted. Label-free protein quantification was based on the Minora algorithm of PD2.4 using the precursor abundance based on intensity and a signal-to-noise ratio >5. Normalization was performed by using the total peptide amount method. Imputation of missing quan values was applied by using abundance values of 75% of the lowest abundance identified per sample. For the reference proteome analysis used for master protein abundance correction of the phosphoproteome data, phosphopeptides were excluded from quantification. Differential protein and phosphopeptide abundance was defined as a fold change of >2, ratio-adjusted pvalue <0.05 (pvalue/log4ratio) and at least identified in 2 of 3 replicates of the sample group with the highest abundance.

### ROS and [Ca^2+^]_cyt_ measurements

Seedlings were grown vertically on Hoagland agar medium (Hoagland’s No. 2 Basal Salt Mixture; Sigma-Aldrich, Germany) for 16 days before harvesting the leaf discs (about 1 mm in diameter), or approximately 70% of the roots for ROS and [Ca^2+^]_cyt_ measurements (Vadassery et al., 2009; Vadassery and Oelmüller, 2009; Johnson et al., 2011).

For ROS measurement, root tissue was incubated in sterile water in a 96-well plate in the dark at room temperature for 1 hour. Prior to the elicitor treatment, water was replaced by 150 μL of assay solution containing 2 μg/mL horseradish peroxidase (Sigma-Aldrich, Germany) and 100 μM luminol (FUJIFILM Wako Pure Chemical Corporation, Japan).

The [Ca^2+^]_cyt_ concentration was inferred from aequorin-based luminescence (Knight et al., 1991). Leaf discs and root tissue were incubated overnight in 150 μL of 7.5 μM coelenterazine solution (P.J.K. GmbH, Germany) in a 96-well plate in the dark at room temperature.

Bioluminescence counts from elicitor application were recorded as relative light units (RLU) with microplate luminometer (Luminoskan Ascent version 2.4, Thermo Electro Corporation, Germany or Mithras LB940, Berthold, Germany).

Cellobiose (C7252, Sigma-Aldrich, Germany), cellotriose (C1167, Sigma-Aldrich, Germany, or 0-CTR-50MG, Megazyme, Ireland) and chitohexaose (OH07433, Carbosynth, United Kingdom) were used as elicitors. Concentration of elicitors, unless specified, is 10 μM for cellotriose and chitohexaose, and 1 mM for cellobiose. All elicitors were dissolved and diluted with distilled water.

### Nucleic acid isolation, PCR and qPCR

Plant tissue was homogenized in liquid nitrogen. DNA extraction was performed according to Doyle (1990). RNA extraction was done with Trizol− reagent (Thermo-Fisher Scientific, Germany), treated with Turbo DNA-free™ Kit (Thermo-Fisher Scientific, Germany), and reverse transcribed with RevertAid Reverse Transcriptase (Thermo-Fisher Scientific, Germany) according to the manufacturer’s instructions.

Genotyping of back-crossed F2 mutant population was achieved by PCRs with genomic DNA. PCRs were run with DreamTaq DNA Polymerase (Thermo Fisher Scientific, Germany) in a thermal cycler (Applied Biosystems SimpliAmp Thermal Cycler, Thermo Fischer Scientific, Germany). Quantitative PCRs (qPCRs) were performed with Dream Taq DNA Polymerase (Thermo-Fisher Scientific, Germany) with the addition of Evagreen® (Biotum, Germany). CFX Connect™ Real-Time PCR Detection System (Bio-Rad, Germany) was used for running and analyzing qPCRs. The expression of genes was normalized to the housekeeping gene encoding a ribosomal protein (RPS; AT1G34030). The resulting ΔCq values were used for statistical analysis. For the confirmation of SNP in the EMS mutant, a primer pair flanking the SNP site was designed, and the region was amplified with Phusion™ High-fidelity DNA polymerase (Thermo-Fisher Scientific, Germany). The PCR product was purified with NucleoSpin Gel and PCR Clean-up kit (Macherey-Nagel, Germany), and sequenced by Eurofins Genomics, Germany. All primers used are listed in Supplementary Table S1.

### Multiple sequence alignment

Amino acid sequences of malectin RLKs and malectin-like RLKs were retrieved from Uniprot database and aligned with MEGA7 (Kumar et al., 2016) using default Clustal W algorithm (Thompson et al., 1994). The aligned sequences were edited for presentation using BioEdit v. 7.2.5 (http://www.mbio.ncsu.edu/BioEdit/bioedit.html). Accession number for all sequences are listed in Supplementary Table S2.

### Plasmid construction

Full length coding regions of *ARF1* and of *CORK1* (AT1G56145.1) were amplified from the reverse-transcribed RNA (cDNA) using Phusion™ High-fidelity DNA polymerase (Thermo-Fisher Scientific, Germany). The fragments were cloned into entry vector pENTR™/D-TOPO™, and transferred to pB7FWG2.0 destination vector (Karimi et al., 2002) with Gateway™ LR Clonase™ II (Thermo-Fisher Scientific, Germany). Site-directed mutagenesis was carried out to specifically mutate the amino acid residues of interest.

For the kinase activity assay, the cytoplasmic domain of CORK1 (residues 654-1039; CORK1^KD^) was cloned and ligated into the expression vector pET28a using restriction enzymes *Bam*HI and *Eco*RI. Two stop codons were added before the *Eco*RI restriction site, generating a 6X His-Tagged protein at the N-terminus. The mutated form (CORK1^KD-G748E^) was obtained by site-directed mutagenesis. The mutated PCR fragment for the kinase domain was cloned and ligated into the expression vector pGEX1λT using restriction enzymes *Bam*HI and *Eco*RI, generating a glutathione S-transferase (GST) fusion protein at the N-terminus.

To generate the luciferase reporter constructs with the *WRKY30* and *WRKY40* promoters, 2 kb-DNA fragments upstream of the respective start codons were cloned, and ligated into the pJS plasmid (Yoo et al., 2007) using *Nco*I and *Bam*HI restriction sites.

For every construct, the insert sequence was confirmed by Sanger sequencing (Eurofins Genomics). Primers used are listed in Supplementary Table S1.

### Protein Expression, Extraction, Purification and Kinase assay

pET28a vector with CORK1^KD^ and pGEX1λT vector with CORK1^KD-G748E^ were transformed into *E. coli* strain BL21(DE3) pLysS (Novagen). For the expression of CORK1^KD^, the transformed bacteria were grown directly in LB broth (Bertani, 1951) with 34 μg/mL chloramphenicol and 50 μg/mL kanamycin at 37°C for 16 hours with shaking. IPTG (isopropyl β- D-1-thiogalactopyranoside; Carl Roth, Germany) was added to the culture to a final concentration of 1 mM to induce protein expression for 3 hours. For the expression of GST-CORK1^KD-G748E^, the overnight culture was inoculated into LB broth with 34 μg/mL chloramphenicol and 100 μg/mL ampicillin at 37°C with shaking. After O.D._600nm_ reached 0.6, IPTG was added to the broth to final concentration of 1 mM to induce protein expression for 3 hours at 25°C. Cells were collected by centrifugation for 10 minutes at 4°C, 5000 rpm.

Bacterial pellet for CORK1^KD^ was resuspended in extraction buffer containing 50 mM Tris-HCl, pH 8.0, 300 mM NaCl and 0.1% (w/v) CHAPS (3-[(3-cholamidopropyl)-dimethylammonio]-1-propansulfonate hydrate, Sigma-Aldrich, Germany). Sonication was applied to lyse the cells. Cell debris was pelleted by centrifugation for 10 minutes at 4°C, 12000 rpm. Cell lysate was incubated with ProBond Ni-NTA resin (Thermo-Fisher Scientific, Germany) for 0.5 – 1 hour. The resin was washed in the same buffer with 20 mM imidazole for 3 times to remove unbound protein. Finally, His-tagged protein was eluted with the same buffer containing 250 mM imidazole. For GST-CORK1^KD-G748E^, purification was done with Pierce− GST Spin Purification Kit following manufacturer’s instruction. Purified proteins were concentrated and buffer-exchanged in kinase assay buffer (25 mM Tris-HCl, pH 7.5, 10 mM MgCl_2_) using Vivaspin^®□^ 20 ultrafiltration unit (3000 MWCO, Satorius, Germany).

Kinase activity assay was carried out by mixing 2 μg of CORK1^KD^ or GST-CORK1^KD-G748E^ with 3 μg of myelin basic protein (MBP; Sigma-Aldrich, Germany) in kinase reaction buffer (25 mM Tris-HCl, pH 7.5, 10 mM MgCl_2_, 1 mM DTT and 100 μM ATP). The reaction mixtures were incubated at 30°C for 30 min and terminated by adding SDS-PAGE loading buffer. The proteins were separated by SDS-PAGE. Protein phosphorylation was examined by staining with Pro-Q Diamond phosphoprotein gel stain (Thermo-Fisher Scientific, Germany) following manufacturer’s instructions, or with morin hydrate (Wang et al., 2013), and visualized using AlphaImager HP system (ProteinSimple, San Jose, California, USA). Coomassie blue staining (Roti-Blue, Carl Roth, Germany) was conducted to visualize the total protein.

### Complementation of the COM receptor mutant

*CORK1* or *ARF1* in pB7FWG2.0 vector were transformed into the COM non-responsive EMS mutant EMS71 using the floral dip method with *A. tumefaciens* strain GV3101 (Zhang et al., 2006). Complemented plants were selected on soil using a 0.1% (v/v) BASTA solution 14 and 18 days after sowing.

### Transient expression in *A. thaliana*

Transient co-expression of the pFRK1::luciferase reporter (Yoo et al., 2007) with the receptor expression constructs in mesophyll protoplasts of *A. thaliana* Col-0 wild-type was performed as described (Wang et al., 2016). Luminescence was recorded for up to 5 hours in W5-medium containing 200 µM firefly luciferin (Synchem UG) after overnight incubation for 14 hours and subsequent treatment with CT or control solutions. After the measurements, protoplasts were harvested by centrifugation and denatured in 2 × SDS-PAGE loading buffer. The crude extracts were separated on 8% polyacrylamide gels and transferred to nitrocellulose membranes. Membranes were saturated with 5% milk powder in PBS with 0,05% Tween-20 (PBS-T) followed by immunostaining with anti-GFP antibodies (Torrey Pines Biolabs, 1:5000 in PBS-T) and secondary goat-anti-rabbit antibodies coupled to alkaline phosphatase (Applied Biosystems) using CDP-star as substrate.

### Microscopy

Protoplast of *A. thaliana* were mounted on a glass slide with cover slip for microscopic inspection using Axio Imager.M2 (Zeiss Microscopy GmbH, Germany). The bright field and fluorescent images were recorded with a monochromatic camera Axiocam 503 mono (Zeiss Microscopy GmbH, Germany). Digital images were processed with the ZEN software (Zeiss Microscopy GmbH, Germany).

### Statistical tests

Statistical tests were performed using R studio version 1.1.463 with R version 4.1.2. Figures were plotted using Python 3.7.4 and arranged with LibreOffice Draw 5.1.6.2.

### Data availability

Raw sequences for the GWAS have been deposited in the Gene Expression Omnibus (GEO) database (accession no. GSE197891). For transcriptome analysis, raw sequences and the count tables after DESeq2 analysis have been deposited in the Gene Expression Omnibus (GEO) database (accession no. GSE198092). Lists of differentially expressed genes mentioned here are provided in Supplementary Dataset S1. The mass spectrometry proteomics data have been deposited to the ProteomeXchange Consortium via the PRIDE (Perez-Riverol et al., 2022) partner repository with the dataset identifier PXD033224. Lists of significantly changed phosphopeptides mentioned here are provided in Supplementary Dataset S2.

## Results

### Identification of the CelloOligomer Receptor Kinase 1 (CORK1)

To identify proteins involved in COM perception, an ethyl methanesulfonate (EMS)-treated seedling population generated from the wild-type pMAQ2 aequorin line (AeqWT) was screened. Roots from individual F2 seedlings were used to monitor [Ca^2+^]_cyt_ elevation upon 1 mM cellobiose (CB) application. One mutant (designated as EMS71) showed no [Ca^2+^]_cyt_ elevation in response to CB (1 mM) and CT (10 μM). The non-responsive phenotype was confirmed in the F3 generation in both root and leaf tissues (Fig. 1A and 1B). Since [Ca^2+^]_cyt_ elevation induced by chitin was not affected (Fig. 1C), EMS71 is specifically impaired in COM perception.

**Figure 1.**
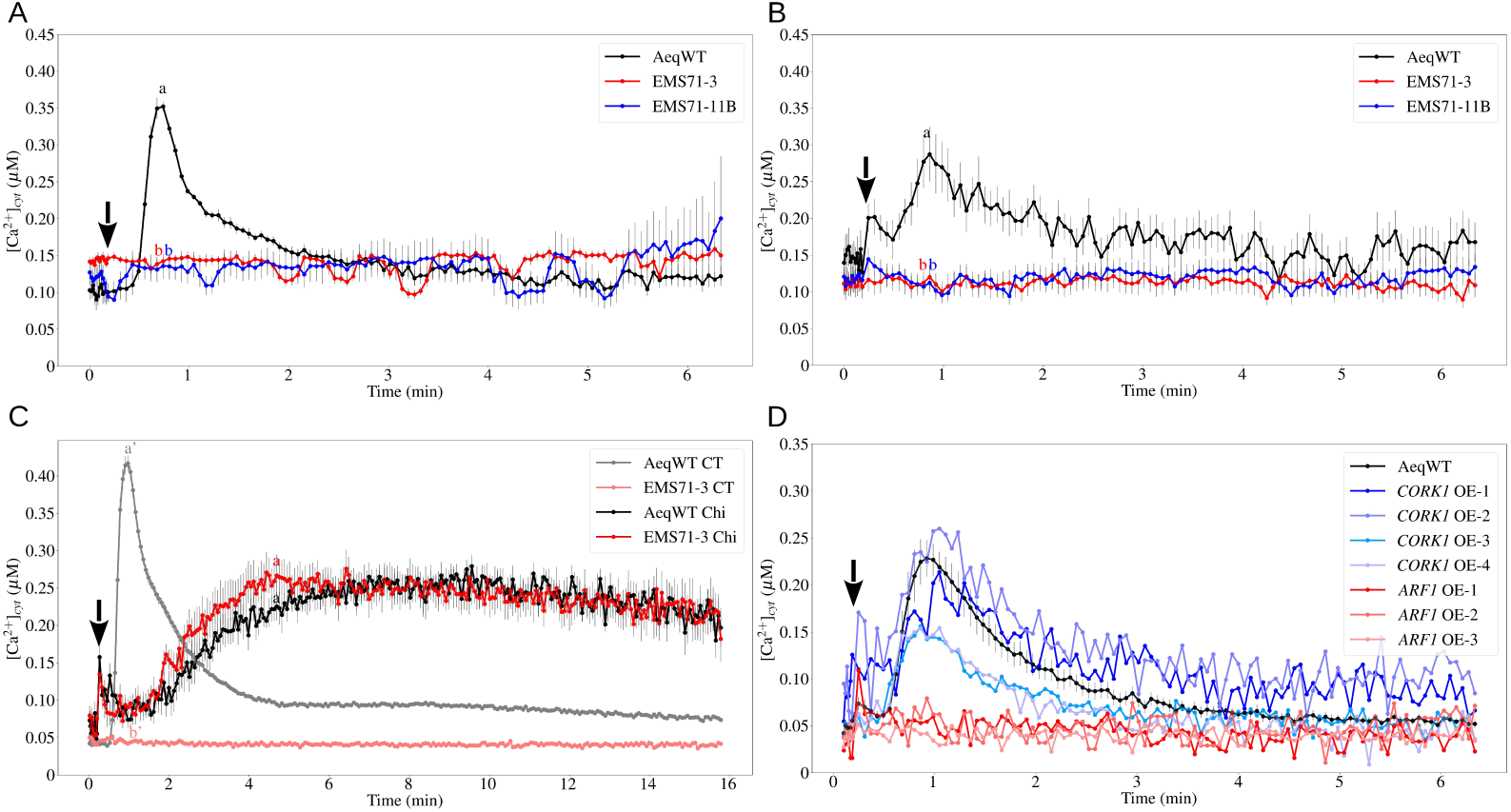
Identification of CORK1 as COM receptor through EMS mutagenesis. A and B, Cytoplasmic m elevation by 10 µM CT in root (A) and leaf (B) tissues. Error bars represent SEs from at least 10 gs. C, Cytoplasmic calcium elevation by 10 µM CT or 10 µM chitohexaose (Chi) in root tissue. Error present SEs from 8 seedlings. D, Cytoplasmic calcium elevation by 10 µM CT in leaf tissue of EMS complemented with *CORK1* or *ARF1*. Error bars represent SEs from 12 seedlings for aequorin wild-AeqWT). Arrows indicate the onset of elicitor application. Statistical significance at the peak value termined by Tukey’s HSD test with P < 0.05, and is indicated by different lower-case letters. All ments were repeated at least 3 times with similar results.

EMS71 was back-crossed to AeqWT, and the F2 population was divided into responsive and non-responsive groups. DNA from these two groups were extracted, sequenced and analyzed as described in Materials and Methods. Finally, two candidate genes, At1g59750 (*ARF1*; A248V) and At1g56145 (*CORK1*; G748E), were selected for further confirmation.

The two candidate genes were over-expressed with CaMV 35S promoter in EMS71. Upon CT application, [Ca^2+^]_cyt_ elevation was only detected in EMS71 seedlings transformed with *CORK1* construct, but not in those with *ARF1*, suggesting *CORK1* is responsible for the [Ca^2+^]_cyt_ elevation induced by COMs (Fig. 1D).

In addition, two T-DNA insertion mutant lines, *cork1-1* and *cork1-2*, were crossed to AeqWT for [Ca^2+^]_cyt_ measurements. In the F2 generation, ∼25% of the seedlings were non-responsive to CT (Fig. 2A). Genotyping showed that the responsive seedlings were either segregated wild-type (SWT) or heterozygotes, while non-responsive seedlings were all homozygous (HO) for the T-DNA insertion (Fig. 2B). The *CORK1* transcript level was significantly reduced in the HO seedlings compared to SWT seedlings (Fig. 2C). This demonstrates that CORK1 is required for COM perception in *Arabidopsis*.

**Figure 2.**
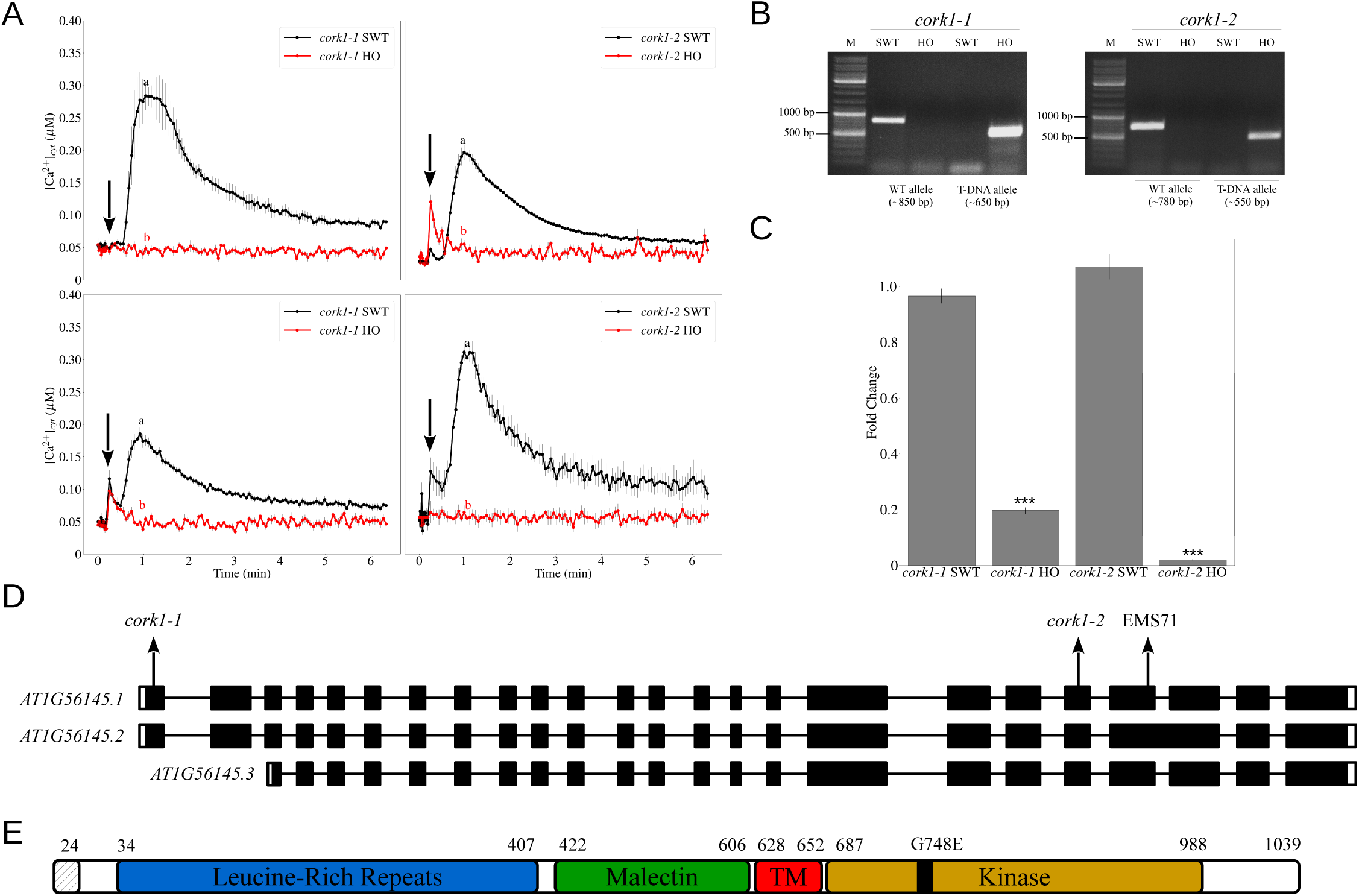
T-DNA mutants for *CORK1* do not respond to COM. A, Cytoplasmic calcium elevation by 10 µM CT in root (upper panels) and leaf (bottom panels) of T-DNA mutants crossed to aequorin wild-type. Error bars represent SEs from at least 5 seedlings. SWT/HO: segregated wild-type/homozygous mutant e cross to aequorin wild-type. Arrows indicate the onset of elicitor application. Statistical significance at the peak value was determined by Tukey’s HSD h P < 0.05, and is indicated by different lower-case letters. The experiment was repeated at least 3 times with similar results. B, Genotyping of the SWT seedlings. Wild-type allele is confirmed with the primer set LP and RP of the respective T-DNA insertion line. T-DNA allele is confirmed with the set LB_SALK and RP of the respective T-DNA insertion line. Annealing temperature for the PCR reactions is 58 °C. M: DNA marker (ladder); bp: base *CORK1* expression in root tissue of SWT and HO seedlings. Error bars represent SEs from 3 independent biological replicates, each with 5 seedlings. al significance was determined by Student’s T-test based on ΔCq values between the two genotypes (***P < 0.001). D, Gene model for *CORK1* 6145). Two T-DNA insertion mutants used in this study are named as *cork1-1* (SALK_099436C; N671776) and *cork1-2* (SALK_021490C; N674063). of the SNP induced by EMS mutagenesis is labeled as EMS71. Arrows indicate the approximate location of T-DNA insertions and SNP on the gene. E, ed protein structure of CORK1. Positions of amino acid residues are shown in numbers. The first 24 amino acids are predicted to be a signal peptide. indicates the amino acid substitution from glycine to glutamate found in EMS71. TM: transmembrane domain.

The gene model for *CORK1* predicts three RNA isoforms (Fig. 2D). At1g56145.1 (lacking an intron near the 3’ end), At1g56145.2 (deduced from the complete DNA sequence) and At1g56145.3 (omitting the first 309 nucleotides from the 5’ end). In the *cork1-1* and *cork1-2* mutants, the T-DNAs were inserted in the first exon and the exon located near the 3’ end, respectively (Fig. 2D). The SNP for EMS71 is caused by a G→A exchange of the 2243^rd^ nucleotide, converting the 748^th^ glycine residue to a glutamic acid (Fig. 2D and 2E). Thus, the latter mutation affects all three predicted RNA isoforms.

Based on the sequence for the first isoform, CORK1 is annotated as a leucine-rich repeat transmembrane protein kinase, with a predicted 24-amino acid long signal peptide at the N-terminus, followed by leucine-rich repeat (LRR) domains, and a malectin domain (MD). After the transmembrane domain, a Ser-Thr/Tyr kinase domain is predicted to reside in the cytoplasm (Fig. 2E).

### *CORK1* encodes a functional receptor kinase

To determine whether *CORK1* encodes a LRR receptor kinase, subcellular localization was first examined by transfecting *A. thaliana* protoplasts with a 35S::CORK1-GFP construct. The GFP signal at the plasma membrane confirmed that CORK1 is a membrane-associated protein (Fig. 3A).

**FIGURE 3.**
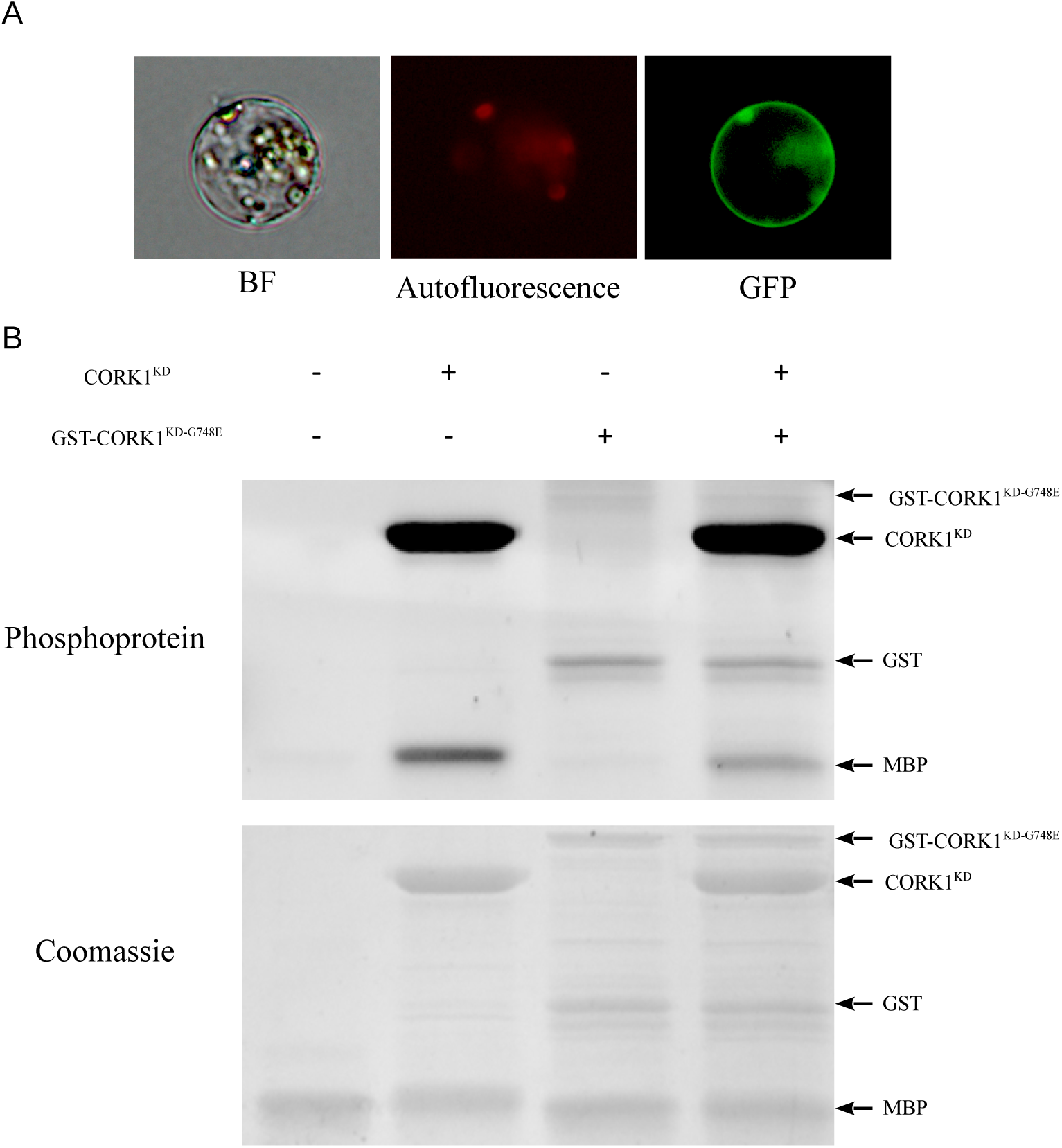
*CORK1* encodes a functional membrane-bound receptor kinase. A, Subcellular localization of gged CORK1 in *Arabidopsis* mesophyll protoplast. BF: bright-field. B, Phosphorylation of the e MBP (myelin basic protein) by CORK1^KD^ but not by CORK1^KD-G748E^.

Next, the cytoplasmic region encompassing the kinase domain (CORK1^KD^) was cloned into the expression vector pET28a to characterize its kinase activity. GST-CORK1^KD-G748E^ was also constructed into pGEX1λT to test whether the mutation found in the kinase domain of EMS71 affects the kinase activity. Figure 3B shows that the substrate myelin protein bovine (MBP) was only phosphorylated by CORK1^KD^ but not by the mutated form GST-CORK1^KD-G748E^. At the same time, CORK1^KD^ exhibited strong autophosphorylation. This suggests that *CORK1* encodes a functional kinase domain, and the G748E mutation disrupted the kinase activity.

### *CORK1* mutant failed to produce ROS upon COM perception

Besides [Ca^2+^]_cyt_ elevation, COMs also induce ROS production, albeit less than classical PAMPs like chitin (Johnson et al., 2018). In SWT roots, but not those in HO, of *cork1-1* and *cork1-2* seedlings, ROS was produced after CT treatment. ROS production was detected upon chitin treatment in both SWT and HO (Fig. 4A and 4B). This suggests that CORK1 is required for COM-, but not chitin-induced ROS production.

**FIGURE 4.**
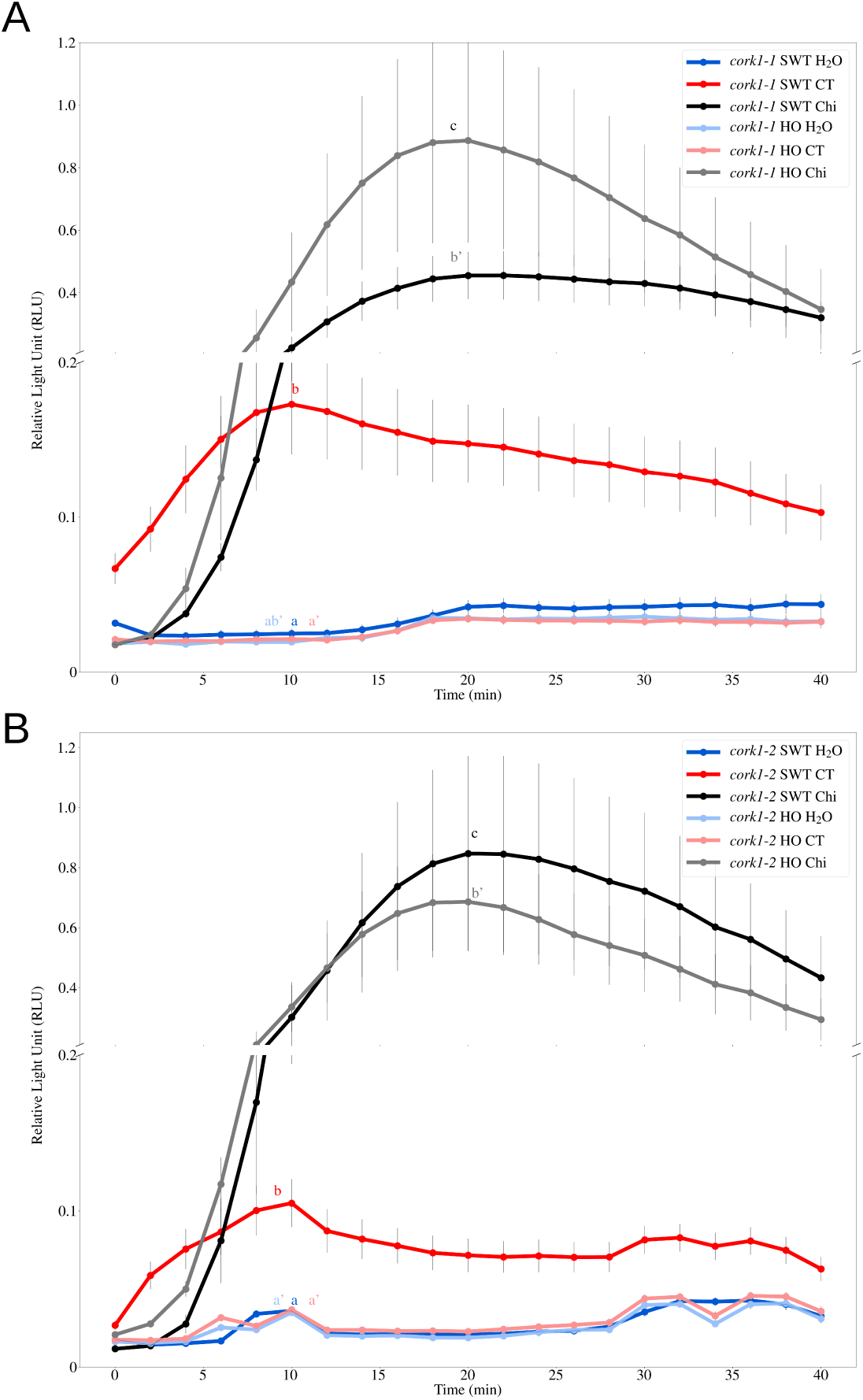
*cork1* mutants failed to induce ROS production upon COM perception. A and B, CT (10 µM) ROS production in root tissue in SWT but not in HO seedlings of *cork1-1* (A) and *cork1-2* (B). ROS ion by application of 10 µM chitohexaose (Chi) was not affected by the mutation. SWT/HO: ted wild-type/homozygous mutant from the cross to aequorin wild-type. Error bars represent SEs least 6 seedlings for each treatment. Statistical significance at the peak value was determined by HSD test with P < 0.05, and is indicated by different lower-case letters. The experiment was repeated 3 times with similar results.

### Up-regulation of *WRKY30* and *WRKY40* mRNA level by COMs is *CORK1*-dependent

Since CB activates *WRKY30* and *WRKY40* expression (Souza et al., 2017; Johnson et al., 2018), we checked whether the activation of these genes requires *CORK1*. CT and chitin were applied to roots of SWT and HO seedlings of *cork1-1* and *cork1-2*. After 1 hour, the *WRKY30* and *WRKY40* transcript levels in SWT of both T-DNA lines were up-regulated ∼30- and ∼15-fold, respectively (Fig. 5A and 5B). On the other hand, no significant response to CT was observed in the HO mutants (Fig. 5A and 5B). Chitin stimulated the *WRKY30* transcript level ∼10-fold and that of the WRKY40 ∼2-fold in both genotypes (Fig. 5A and 5B). This demonstrates COM-mediated activation of *WRKY30* and *WRKY40* requires CORK1.

**FIGURE 5.**
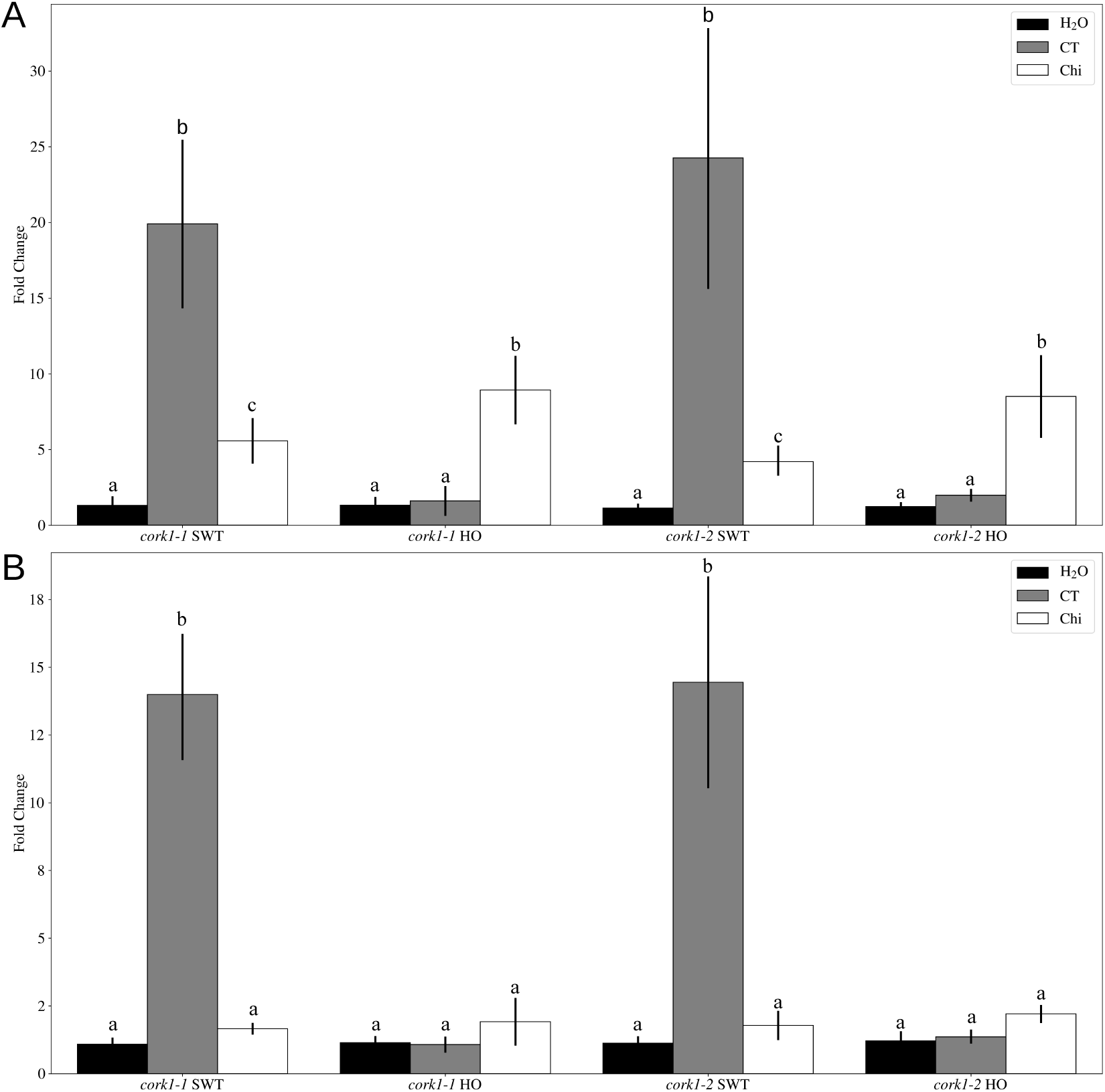
Up-regulation of *WRKY30* and *WRKY40* mRNA level in root tissue by COM is *CORK1*-ent. A and B, *WRKY30* (A) and *WRKY40* (B) mRNA level 1 h after 10 µM CT or 10 µM chitohexaose eatment in *cork1-1* and *cork1-2* SWT (segregated wild-type) and HO (homozygous mutant) from the aequorin wild-type. Values were normalized to water treatment on the same genotype. Error bars nt SEs from at least 4 independent biological replicates, each with 16 seedlings. Statistical ance was determined by Tukey’s HSD test based on ΔCq values with p-value < 0.05, and indicated rent lower-case letters.

### Two Phe residues in the malectin domain are important for CT response

Sequence alignment of the *Arabidopsis* LRR-MD RLKs demonstrated that two Phe residues within the MD (F520 and F539) are highly conserved in all MD RLKs (Fig. 6A and Supplementary Fig. S1) and Malectin-Like (MLD) RLKs (Supplementary Fig. S2). It has been suggested that aromatic rings of amino acids interact with the apolar side of carbohydrate (Boraston et al., 2004; Pires et al., 2004; Schallus et al., 2010). Therefore, we changed the two conserved Phe residues to Ala. In the EMS71 mutant transformed with a 35S::CORK1-GFP construct, the [Ca^2+^]_cyt_ elevation in response to CT application was restored. The [Ca^2+^]_cyt_ response in plants transformed with either of the two Phe mutant versions (F520A or F539A) was significantly reduced, and no [Ca^2+^]_cyt_ could be observed in plants transformed with the double mutated version (Fig. 6B). To further support the importance of the two Phe residues, mesophyll protoplasts of *A. thaliana* were co-transformed with the pFRK1::luciferase reporter and either the wild-type or the double-mutated version of CORK1. The co-expression of wild-type CORK1 with the reporter gene conferred responsiveness to the treatment with 1 µM CT, which was absent when the mutated form was expressed (Fig. 6C - 6F). This suggests that the two conserved Phe residues are important in COM perception in *Arabidopsis*.

**Figure 6.**
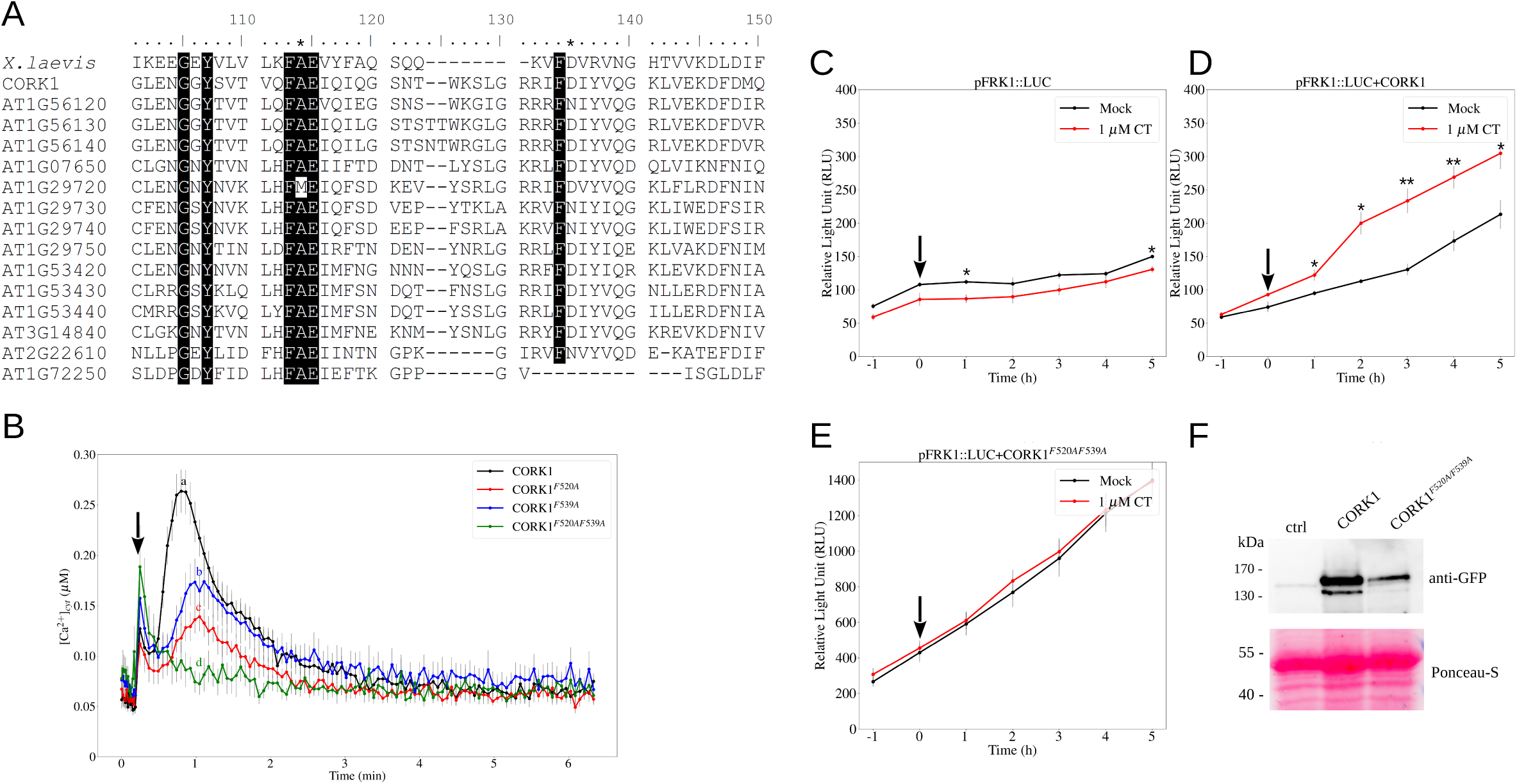
Two conserved phenylalanine residues in the malectin domain are important for COM perception. A, Alignment of the amino acid sequences from n of *X. laevis* and the malectin domains present in *A. thaliana* LRR-malectin RLKs. Shown here are amino acids from position 101-150 of the alignment. hade indicates conserved amino acid residues over 90% threshold. The two conserved phenylalanine residues are indicated with an asterisk. B, smic calcium elevation by 10 µM CT in leaf tissue of EMS71 complemented with CORK1, or with single (CORK1^F520A^/CORK1^F539A^) or double 1^F520AF539A^) mutation in the two conserved phenylalanine residues. Error bars represent SEs from 12 seedlings. Arrow indicates the onset of elicitor ion. Statistical significance at the peak value was determined by Tukey’s HSD test with P < 0.05, and is indicated by different lower-case letters. The ent was repeated 3 times with similar results. C-E, Protoplasts from *A. thaliana* Col-0 were transfected with the pFRK1::Luciferase (pFRK1::LUC) construct (C), or the reporter construct plus the CORK1 receptor (D), or the reporter construct plus the double mutated version CORK1 ^F520AF539A^ (E). show luciferin-dependent light emission over time after treatment with water (Mock) or 1 µM CT. Arrow indicates the onset of elicitor application at 0 data point represents the mean value from 4 technical replicates. Error bars represent SEs. Statistical significance was determined by Student’s T-test n the two treatments (*P < 0.05; **P < 0.01). The experiment was repeated 4 times with similar results. F, Western blot indicating the expression of both versions of GFP-tagged CORK1. ctrl: control, protoplast transfected with pFRK1::LUC reporter construct only.

### Transcriptome analysis uncovered COM/CORK target genes

To identify the biological functions of COMs, we performed transcriptome analysis with the roots of *cork1-2* SWT and HO seedlings 1 hour after the application of either 10 μM CT or water. Among the 23106 mapped genes, 561 genes were up- and 54 genes down-regulated by CT in a *CORK1*-dependent manner. On the contrary, only 2 genes were significantly up-regulated and no genes were down-regulated by CT in HO (Fig. 7A; Supplementary Dataset S1). This shows the high specificity of *CORK1* to COMs.

**Figure 7.**
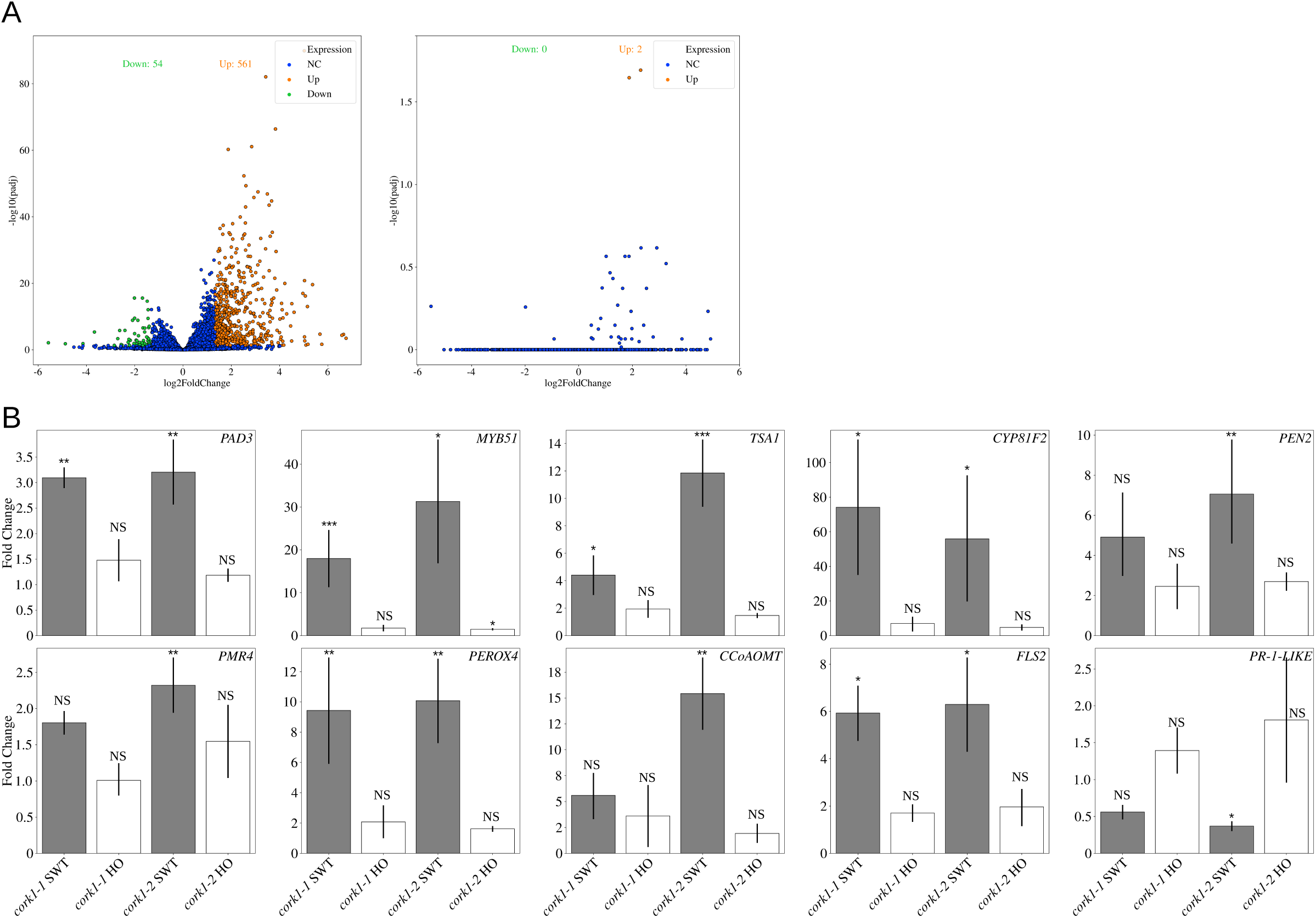
CT-regulated genes. A, Volcano plots showing the distribution of differentially expressed genes in root tissue. Left: 10 µM CT treatment compared to ontrol in SWT. Right: 10 µM CT treatment compared to water control in HO. NC: no change; Up: up-regulation; Down: down-regulation; padj: adjusted using Benjamini and Hochberg method. The FDR cutoff value is set as 0.1. B, qPCR analysis of candidate genes regulated by 10 µM CT in root tissue T and HO seedlings. Values were normalized to water treatment on the same genotype. SWT/HO: segregated wild-type/homozygous mutant from the f *cork1-2* to aequorin wild-type. Error bars represent SEs from 4 independent biological replicates, each with 16 seedlings. Statistical significance was ned by Student’s T-test based on ΔCq values (NS: not significant; *P < 0.05; **P < 0.01; ***P < 0.001).

Gene ontology (GO) enrichment analysis showed a profound increase in genes involved in tryptophan biosynthesis, cell wall modification and secondary metabolite production (Supplementary Fig. S3). The genes for ASA1 (anthranilate synthase α subunit 1) and ASB1 (anthranilate synthase β subunit 1), which carry out the first step in the tryptophan biosynthesis from chorismate, were ∼10 fold and for TSA1 (tryptophan synthase α chain), which catalyzes the last step in the biosynthesis, ∼8 fold up-regulated by CT (Table 1 and Fig. 7B).

**TABLE 1.**
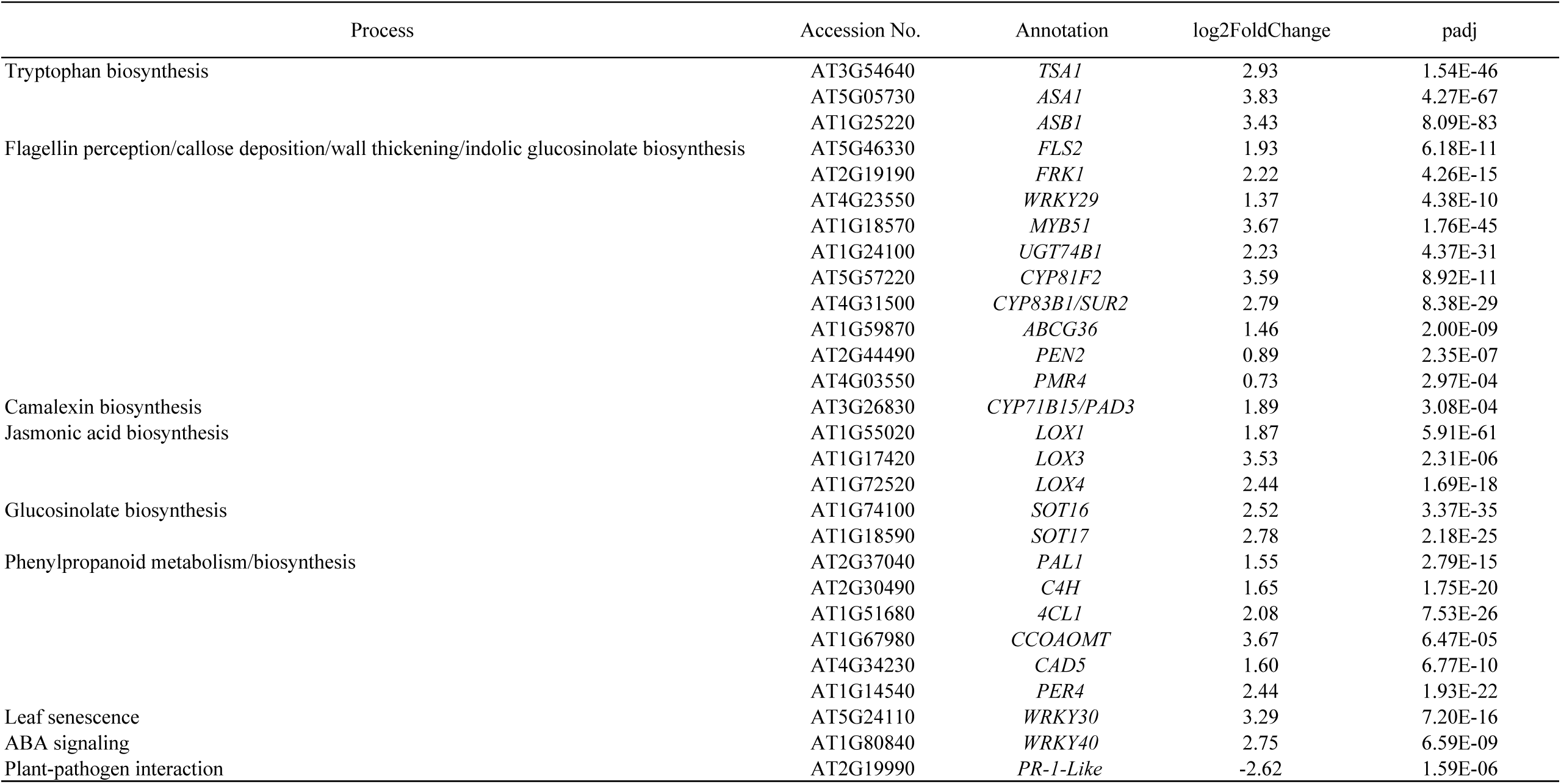
Differentially expressed candidate genes by 10 µM CT compared to water control in root tissue of *cork1-2* segregated wild-type from the cross to aequorin wild-type. Process Accession No. Annotation log2FoldChange padj

Among the first 15 categories for the most strongly regulated genes, 5 categories are related to “cell wall” functions. All of them center around callose deposition and cell wall thickening, and most of these genes/proteins are described in the context of defense (Supplementary Fig. S3). Genes in these categories include *FLS2* (∼4 fold), *MYB51* (∼13 fold), *UDP-glycosyltransferase 74B1* (∼4.5 fold), the cytochrome P450 enzymes *CYP81F2* (∼12 fold) and *CYP83B1* (∼7 fold) as well as the ABC-transporter gene *ABCG36* (∼3 fold). Similarly, genes involved in lignin biosynthesis (phenylpropanoid metabolism) were also up-regulated, such as those for cinnamate-4-hydroxylase (C4H, ∼3 fold), 4-coumarate-CoA ligase 1 (4CL1, ∼4 fold), phenylalanine ammonia lyase 1 (PAL1, ∼3 fold), and for three enzymes important for lignin production, Caffeoyl-CoA 3-O-methyltransferase (CCoAOMT, ∼13 fold), cinnamyl alcohol dehydrogenase 5 (CAD5, ∼3 fold) and peroxidase 4 (PER4, ∼5 fold; Fernández-Pérez et al., 2015; Barros et al., 2019). *PEN2* and *PMR4/GSL5*, encoding a myrosinase and a callose synthase, respectively, were only slightly up-regulated (∼1.7 fold; Table 1 and Fig. 7B).

In addition to cell wall-related genes, *SOT16* and *SOT17*, which encode sulfotransferases for glucosinolate production, were up-regulated ∼5.5 fold. Likewise, the transcript level for *CYP71B15* (*PAD3*), which is required for camalexin production, was ∼3.5-fold up-regulated. In line with qPCR analyses and previous reports (Souza et al., 2017; Johnson et al., 2018), *WRKY30, WRKY40* and the lipoxygenase genes involved in jasmonic acid synthesis *LOX1* (∼3.5 fold), *LOX3* (∼11.5 fold) and *LOX4* (∼5.5 fold) responded to COMs. Finally, genes for proteins in the FLS2 signaling pathway were up-regulated, such as *FRK1* (∼4.5 fold) and *WRKY29* (∼2.5 fold; Asai et al., 2002; Table 1 and Fig. 7B).

Interestingly, most of the down-regulated genes are involved in ion homeostasis, and a *PR-1 like* gene which mediates defense (At2g19990, ∼0.16 fold; Fig. 7B and Supplementary Fig. S4).

In summary, COM perception by CORK1 induces cell wall reinforcement, defense-related secondary metabolite synthesis, and crosstalk with other signaling components.

### COM/CORK-mediated changes in the phosphoproteome pattern in roots

To identify early COM/CORK1 targets, the phosphoproteomes of SWT and HO roots were analyzed 5 and 15 minutes after 10 μM CT or water (control) application. Most of the proteins with a significantly altered phosphorylation state are related to (i) cellulose synthase complex (CSC) functions and translocation to the plasma membrane, (ii) the ER secretory pathway and protein sorting, (iii) proteins involved in signal transduction, or (iv) defense/stress responses (Supplementary Fig. S5 and Supplementary Dataset S2).

Cellulose synthases 1 and 3 (CESA1, -3) of the CSC are required for cellulose synthesis for the primary cell wall and the protein is rapidly phosphorylated at Ser24 and Ser176, respectively, in response to CT application. Mutations of CESAs phosphorylation sites modulate anisotropic cell expansion and bidirectional mobility of the cellulose synthase (Chen et al., 2010). Besides CESAs, we also identified Cellulose Synthase-Interactive 1 (CSI1) as a phosphorylation target of CT at Thr37. Association of CSC with cortical microtubules is mediated by CSI1 and the protein contains multiple phosphorylation sites potentially involved in regulatory processes (Jones et al., 2016). Loss of function CSI1 mutants are impaired in the dissociation of the CSC from the microtubules during their passage to the plasma membrane which results in cellulose deficiency in the mutant cell walls (Li et al., 2015). The COMPANION OF CELLULOSE SYNTHASE 1 and 2 (CC1/CC2) and the N-terminal domain in CSI are responsible for the connection of the CSCs to the cortical microtubules and *csi1* mutants are impaired in microtubule stability under salt stress (Endler et al., 2015; Speicher et al., 2018), but also for CSC delivery to the plasma membrane and its recycling (Lei et al., 2012; Lei et al., 2015). In addition to its role in trafficking and mobility of CSCs, microtubules also influence the orientation and crystallinity of cellulose (Lei et al., 2015). Thus, CESA1 and CSI1 are two central players in cellulose repair mechanism, and are phosphorylated by COM/CORK.

Among the proteins involved in the endomembrane system and the secretory pathway are two GTPases which regulate membrane trafficking: AGD5, a GTPase-activating protein operating at the *trans*-Golgi network (Stefano et al., 2010) and RABA5C, a GTPase that specifies a membrane trafficking pathway to geometric edges of lateral root cells (Kirchhelle et al., 2019). The 1-phosphatidylinositol-3-phosphate 5-kinase is involved in maintenance of endomembrane homeostasis including endocytosis and vacuole formation (Hirano et al., 2011); the exocyst complex component SEC8 participates in the docking of exocytic vesicles with fusion sites on the plasma membrane and the formation of new primary cell wall; the vacuolar sorting protein 41 regulates vacuolar vesicle fusions and protein sorting together with phosphoinositides (Brillada et al., 2018); a SNARE protein as part of a complex facilitates trafficking in the endomembrane system including distinct secretory and vacuolar trafficking steps. The phosphorylated myosin mediates the organization of actin filament and vesicle transport along the filaments. Finally, MAPK 17 influences the number and cellular distribution of peroxisomes through the cytoskeleton-peroxisome connection (Frick and Strader, 2018). Not surprisingly, stimulation of membrane trafficking, protein sorting and secretion also affects enzymes involved in cellulose, callose and other polysaccharide biosynthesis, such as CESAs, CSI1, callose synthase and a regulator of callose deposition, the PAMP-Induced coiled coil protein At2g32240 (Wang et al., 2019). The cytosolic UDP-glucuronic acid decarboxylases, At3g46440 and At5g59290, produce UDP-xylose, which is a substrate for many cell wall carbohydrates including hemicellulose and pectin. UDP-xylose is also known to feedback regulate several cell wall biosynthetic enzymes, many of them are associated with the endomembrane system (Pattathil et al., 2005; Kuang et al., 2016; Zhong et al., 2017). Phosphorylated proteins with related functions control auxin translocation at the plasma membrane (At1g56220, ABCG36).

Inspection of target proteins at different time points uncovered that the canonical immunity-related mitogen-activated protein kinases MPK3 and MPK6 showed increased phosphorylation (Ser16 of MPK3 and Tyr223 of MPK6) at both time points after CT application in SWT roots. Likewise, the calmodulin-binding protein IQM4 (at Ser505, Ser509, Ser520 and Ser525) and the Ca^2+^-dependent protein kinase CPK9 (at Ser69) are among the most phosphorylated targets at both time points and link stress-induced Ca^2+^ signaling to abscisic acid (Zhou et al., 2018; Chen et al., 2019). On the other hand, phosphorylation of the plasma membrane-localized FERONIA (at Ser695), SERK1 (at Thr450 and Thr463) and MAPKKK3 which mediates MPK3/6 activation by at least four pattern-recognition receptors (FLS2, EFR, CERK1, and PEPRs; Bi et al., 2018) was only detectable at the early time point. On the other hand, phosphorylation at Ser716 of the Tumor Necrosis Factor Receptor-Associated Factor (TRAF) 1B, which involved in immune receptor turnover, was increased 5 minutes but decreased 15 minutes after CT treatment (Huang et al., 2016; Table 2).

**TABLE 2.**
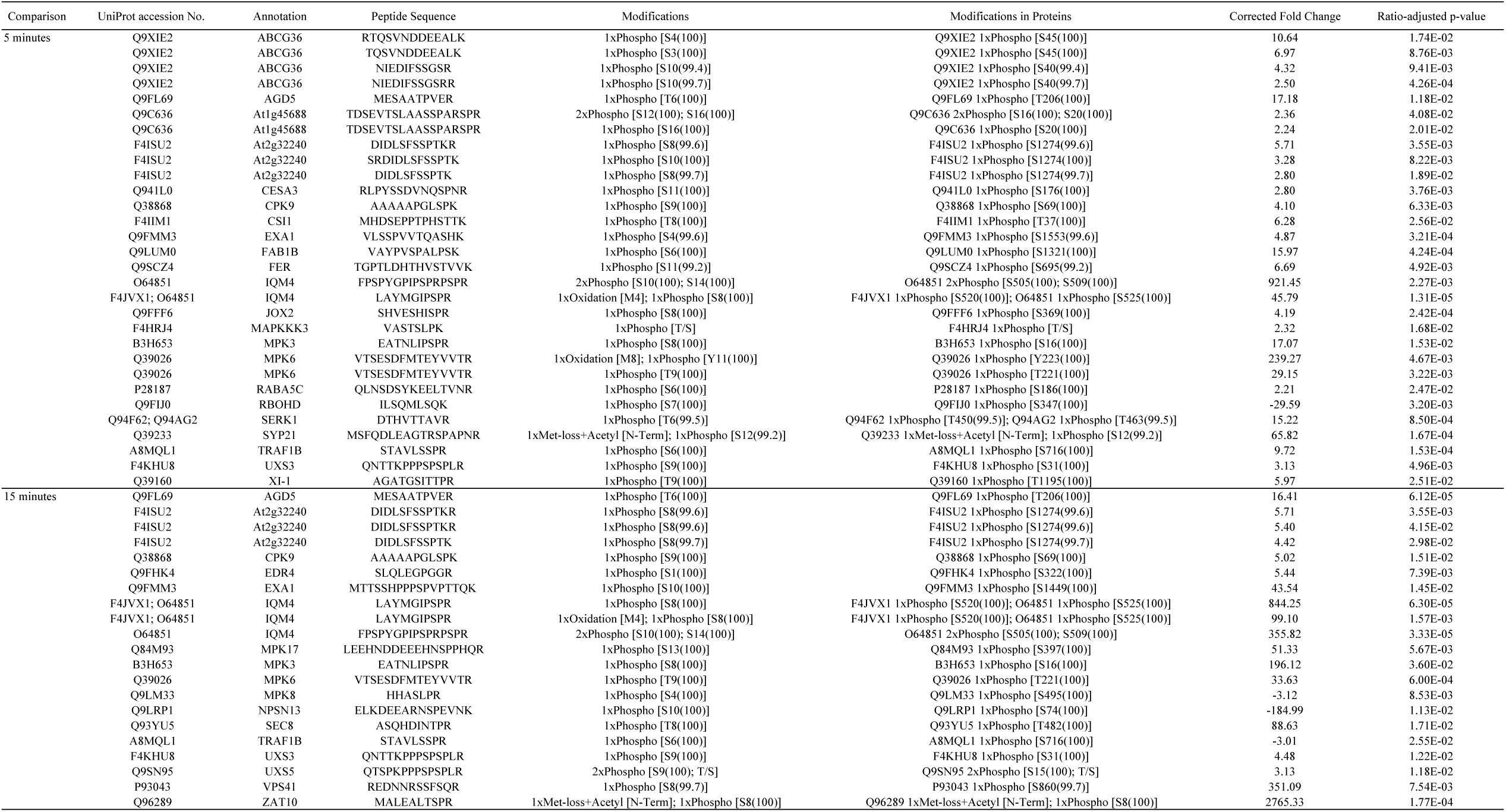
Candidate proteins with significant alteration in phosphorylation upon 10 μM CT treatment compared to water control in root tissue of *cork1-2* SWT. Numbers in the parentheses indicate the probability of the modification on the amino acid residue. Calculation for the corrected fold change and the ratio-adjusted p-value is described in the Materials and Methods section. SWT: segregated wild-type from the cross to aequorin wild-type

Among the defense-related proteins, phosphorylation of RBOHD at Ser347 was significantly decreased at the early time point (Table 2). The MPK8 module negatively regulates ROS accumulation through controlling expression of the *RBOHD* gene and the phosphorylation state decreased 15 min after CT application. Takahashi *et al*. (2011) proposed that Ca²^+^/CaMs and the MAP kinase phosphorylation cascade converge at MPK8 to monitor or maintain ROS homeostasis. Likewise, phosphorylation of JOX2 at Ser369 which catalyzes the hydroxylation of jasmonic acid to 12-OH-jasmonic acid and thus restricts the generation of the active jasmonic acid-isoleucine, was only detectable at the early time point. Also EXA1 (Essential for potexvirus Accumulation 1) controlling virus infection was only phosphorylated at Ser1553 at the early time point (Hashimoto et al., 2016). In contrast, phosphorylation of EDR4 (Enhanced Disease Resistance 4) which represses salicylic-acid-mediated resistance, and ZAT10, a repressor of abiotic stress and jasmonic acid responses, was only stimulated 15 min after CT application (Wu et al., 2015; Xie et al., 2019). The ABC transporter G36 which controls pathogen entry into cells, was phosphorylated at both time points and its mRNA was upregulated in our expression analysis (Table 1). The identified targets of COM/CORK1 signaling show dynamic changes in the protein phosphorylation pattern. Considering the different roles of these proteins in activating or repressing signaling and defense responses, it can be speculated that COM/CORK1 signaling establishes a moderate immune response and maintains its homeostasis.

## Discussion

We demonstrate that CORK1, a leucine-rich repeat-malectin receptor kinase, is required for COM-mediated rapid increase in [Ca^2+^]_cyt_ level and stimulation of ROS production in *Arabidopsis*. Transcriptome analyses uncovered CT-regulated and CORK1-dependent target genes of a proposed COM/CORK1 signaling pathway. Major CT/CORK1 target genes are involved in cell wall strengthening, secondary metabolite metabolism and Trp biosynthesis. Phosphoproteome analysis identified early COM/CORK1 target proteins involved in secretory pathways and vesicle trafficking, the plasma membrane-associated RBOHD, FER and SERK1, MPK3/6, novel MAPKs such as MAPKKK3, MPK17 and MPK8, as well as downstream proteins involved in plant immunity. Interestingly, the different phosphorylation patterns 5 and 15 min after the stimulus and phosphorylation of EXA1, TRAF1B, MPK8, JOX2 and EDR4 demonstrate that CT establishes a balanced defense response. For instance, CT stimulates ROS production. Simultaneously phosphorylation and thus activation of RBOHD was repressed at the early time point (Table 2), and MPK8, which is involved in establishing ROS homeostasis (Takahashi et al., 2011), is phosphorylated. Furthermore, expression of defense-related genes such as *WRKY30*/*40*, is stimulated by COM, while phosphorylation of EXA1, TFAF1B, JOX2 and EDR4 restrict or balance defense responses. A crosstalk to other receptor kinases is demonstrated by FER and SERK1, and potentially the up-regulation of *FLS2* at the mRNA level. The observed downstream responses are consistent with the idea that COM/CORK1 activates processes which maintain cell wall integrity.

### LRR-Malectin receptors are the new players in cell wall surveillance

Malectins were first discovered in *Xenopus* (Schallus et al., 2008). They are located in the ER of animal cells, and bind to diglucosylated N-linked glycans to control glycoprotein quality (Galli et al., 2011; Takeda et al., 2014; Tannous et al., 2015). The ligands of malectins include the disaccharide maltose and nigerose (Schallus et al., 2008; Schallus et al., 2010). The structure of malectins are similar to the carbohydrate-binding modules found in enzymes which degrade the plant cell wall (Schallus et al., 2008; Gilbert et al., 2013; Duan et al., 2016). Although the two Phe residues conserved in all MDs in *A. thaliana* are not conserved in the maltose binding malectin from *Xenopus* (Schallus et al., 2010), the replacement of the two Phe with Ala eliminated CT-induced [Ca^2+^]_cyt_ elevation and reporter gene activation (Fig. 6). This suggests a functional divergent role of malectins in plants and animals. The two Phe residues in the plant MDs might be specifically involved in binding COs with β 1-4-bonds, while non-plant malectins bind maltose and nigerose with α 1-4-bonds.

Besides CORK1, there are 13 additional LRR-MD-RLKs in *Arabidopsis*. They have been shown to be involved in lipopolysaccharide perception (Hussan et al., 2020), pollen tube development (Lee and Goring, 2021) and control of cell death in leaves (Li et al., 2020). Several of them are up-regulated by brassinosteroids and participate in immune responses (Qutob et al., 2006; Hok et al., 2014; Xu et al., 2014). However, whether their MDs bind to sugars is not known. Due to structure and sequence similarity, they might interact with other sugars from cell wall polymers, since the *cork1* phenotype excludes their participation in COM responses.

Intriguingly, the phosphoproteomic study identified FER as a phosphorylation target after CT treatment (Table 2). It harbors a malectin-like domain (MLD), consisting of two tandem malectin domains. It has been shown to regulate CWI and pollen tube development, although known ligands for FER are RALF peptides and pectins (Escobar-Restrepo et al., 2007; Zhang et al., 2020; Tang et al., 2021; Yang et al., 2021b). Together with the phosphorylation of several members of CSC, it is likely that CORK1 controls cell wall repair mechanism, and may coordinate FER upon cell wall damage.

### Crosstalk between CORK1 and other signaling pathways

Activation of *FLS2*, genes involved in FLS2 signaling and FLS2 targets, phosphorylation of MAPKs (MAPKKK3, MPK3 and MPK6), plasma membrane-localized SERK1 and FER by CT/CORK1 suggest crosstalk to other receptor kinases and PAMP-activated defense signaling (Fig. 7, Table 1 and Table 2). Souza *et al*. (2017) and Johnson *et al*. (2018) have shown that combined treatments of CB/CT with either flg22 or chitin trigger higher [Ca^2+^]_cyt_ levels, ROS production and MPK3/6 phosphorylation. CWI signaling and plant defense are tightly coupled, both operate via [Ca^2+^]_cyt_ elevation. Although their Ca^2+^ signatures might differ, the overlap is apparent by the large number of defense-related genes which respond to CT/CORK1 activation and PAMP-induced signaling. Likewise, *WRKY30* and *WRKY40* are downstream targets of COM/CORK1 activation and the transcription factors participate in various biotic and abiotic responses (Scarpeci et al., 2013; Wang et al., 2021). Understanding the crosstalk between CORK1 and other PRRs as well as their signaling components will provide a broader picture of how plants integrate different threats and developmental signals. CWI signaling is also important during many developmental processes, starting from growth, division and differentiation of cells, meristem development, senescence to fertilization. The tissue-specific expression of the different members of the MD-containing receptor kinases might reflect their different roles in monitoring cell wall alterations (Wu et al., 2016).

### CT regulates metabolism of aromatic amino acids and secondary metabolites

Tryptophan-derived secondary metabolites have long been considered as important components for innate immunity, and Trp serves as the starting amino acid for the biosynthesis of camalexin and indolic glucosinolates (Bednarek, 2012). Up-regulation of Trp biosynthesis is crucial for plant defense against fungal pathogens and hemibiotrophs (Ishihara et al., 2008; Consonni et al., 2010; Hiruma et al., 2013).

Although several genes important for flagellin-induced callose deposition (Clay et al., 2009) are also up-regulated by CT, however, callose deposition could not be observed (Souza et al., 2017). This might be due to the low induction of *PEN2* and *PMR4* by CT (Fig. 7). Interestingly, *ABCG36* (*PEN3*) was up-regulated and the gene product, ABCG36/PEN3, was phosphorylated by CT after 5 minutes (Fig. 7 and Table 2). Phosphorylation of the transporter is important for pathogen defense, and it has been proposed that it participates in the deposition of defense-related secondary metabolites (Underwood and Somerville, 2017). Which components can be exported by ABCG36 (PEN3) are not known, but they might as well be important for cell wall repair. Besides differences between COM (DAMP) and flagellin (PAMP), our –omics analyses confirm crosstalks at the signaling levels. Besides COM/CORK1-induced investment into defense, it might play an important role in priming plant immune responses induced by other stimuli.

Another group of CT-stimulated genes involved in phenylpropanoid metabolism (Fig. 7) convert Phe through cinnamic acid (Phe-ammonia lyase), *p*-coumaric acid (cinnamate-4-hydroxylase) to *p*-coumaryol-CoA (4-coumarate:CoA ligase 1). From there, *p*-coumaryol-CoA can be used to synthesize lignin with cinnamyl alcohol dehydrogenase 5 (CAD5), caffeoyl-CoA O-methyltransferases and peroxidase 4 (Fernández-Pérez et al., 2015; Barros et al., 2019). These steps are important in secondary cell wall synthesis.

Apart from aromatic compounds, expression of lipoxygenases involved in jasmonic acid (JA) biosynthesis are up-regulated by CT (Table 1). Moreover, the gene for CML42, a negative regulator of JA signaling and biosynthesis, is down-regulated by CT (Vadassery et al., 2012; Table 1). Furthermore, JOX2, which converts JA to 12-hydroxyjasmonate, an inactive form of JA (Zhang et al., 2021), is phosphorylated at Ser369 in response to CT. The enzyme prevents over-accumulation of JA and its bioactive form JA-Ile under stress (Caarls et al., 2017; Smirnova et al., 2017) and thus, represses basal JA defense responses. These results show that JA is a target of COM signaling. However, it appears that COM treatment established moderate and balanced JA level in the cell. The comparative analysis of the regulated genes and phosphorylation targets suggest that the primary effect of COM/CORK1 signaling is activating cellular processes which strengthen the cell wall and not those promoting cytoplasmic immunity.

This work provides evidence for the first receptor for cellulose breakdown fragments, and demonstrates the importance of the malectin domain for COM sensing. With recent findings of other cell wall breakdown products acting as DAMPs (Claverie et al., 2018; Mélida et al., 2020; Rebaque et al., 2021; Yang et al., 2021a), closer inspection of the other LRR-MD RLK members might be a reasonable strategy to identify receptors for other polysaccharide breakdown products of the cell wall. Our transcriptome and phosphoproteome analyses provide a list of components which are potentially involved in COM signaling, might represent COM targets, or participate in CORK1 crosstalk.

## Acknowledgement

We thank Prof. Dr. Maria Mittag for providing the expression vector pGEX1λT and Prof. Georg Felix for discussions and comments on the work. We also thank Claudia Röppischer, Sarah Mußbach and Christin Weilandt for technical assistance. Special thanks to Yelyzaveta Zhyr for the help on plant transformation and the green house team at Max Planck Institute for Chemical Ecology for taking care of all the *Arabidopsis* transformants. This work was supported by the Deutsche Forschungsgemeinschaft (DFG, German Research Foundation) CRC1127 ChemBioSys (project ID: 239748522) and CRC/TR 124 FungiNet (project Z2; project ID: 210879364).

## Author Contributions

**Y-HT** and **RO** developed the idea and organized the project. **Y-HT**, performed the experiments, collected the samples and data, analyzed the results and plotted the figures. **SSS** generated the EMS mutant population. **JF** performed the protoplast assay and analyzed the data. **TK** carried out the phosphopeptide measurement and data analysis. **AG** assisted on sample collection and protein extraction. **AAB** and **OK** provided the materials and equipment for protein extraction. **Y-HT** and **RO** wrote up the study. All authors contributed to the manuscript.

